# Murine astrotactins 1 and 2 have similar membrane topology and mature via endoproteolytic cleavage catalyzed by signal peptidase

**DOI:** 10.1101/493858

**Authors:** Patricia Lara, Åsa Tellgren-Roth, Hourinaz Behesti, Zachi Horn, Nina Schiller, Karl Enquist, Malin Cammenberg, Amanda Liljenström, Mary E. Hatten, Gunnar von Heijne, IngMarie Nilsson

**Author notes:** To whom correspondence should be addressed: Gunnar von Heijne and IngMarie Nilsson, Department of Biochemistry and Biophysics, Stockholm University, Svante Arrhenius väg 16C, SE-10691 Stockholm, Sweden.

## Abstract

Astrotactins 1 (Astnl) and Astn2 are membrane proteins that function in glial-guided migration, receptor trafficking and synaptic plasticity in the brain, as well as in planar polarity pathways in skin. Here, we used glycosylation mapping and protease-protection approaches to map the topologies of mouse Astn1 and Astn2 in rough microsomal membranes (RMs), and found that Astn2 has a cleaved N-terminal signal peptide (SP), an N-terminal domain located in the lumen of the RMs (topologically equivalent to the extracellular surface in cells), two transmembrane helices (TMHs), and a large C-terminal lumenal domain. We also found that Astn1 has the same topology as Astn2 but we did not observe any evidence of SP cleavage in Astnl. Both Astnl and Astn2 mature through endoproteolytic cleavage in the second TMH; importantly, we identified the endoprotease responsible for the maturation of Astn1 and Astn2 as the endoplasmic reticulum signal peptidase. Differences in the degree of Astn1 and Astn2 maturation possibly contribute to the higher levels of the C-terminal domain of Astn1 detected on neuronal membranes of the central nervous system. These differences may also explain the distinct cellular functions of Astn1 and Astn2, such as in membrane adhesion, receptor trafficking, and planar polarity signaling.

---

Astrotactins are vertebrate-specific integral membrane glycoproteins known to play critical roles in central nervous system (CNS) and skin development (1-4). An understanding of the function of Astn1 and Astn2 in the control of neuronal migration and of synaptic function could be important for treatment of human brain disorders such as epilepsy and autism spectrum disorders. Although the number of gene mutations that can disrupt neuronal migration is large (5), Astn1 is one of a few adhesion receptors shown to directly function in migration (6).

In mouse, there are two astrotactin family members, Astn1 and Astn2 (ASTN1 and ASTN2 in humans). Astn1 is involved in glial-guided neuronal migration early in development (1,3,6,7) through the formation of an asymmetric complex with N-cadherin (CDH2) in the glial membrane (6). Astn2, which is 48% homologous to Astn1 and has two isoforms, is abundant in migrating cerebellar granule neurons where it forms a complex with Astn1, and regulates the trafficking of Astn1 during migration (4). At later stages of development, Astn2 regulates synaptic function by trafficking of other membrane receptors, including the Neuroligins and other synaptic proteins (8). A recent structure of the C-terminal endodomain of Astn2 shows distinctive features responsible for its activity (9). Astn1 and Astn2 are believed to share the same membrane topology, with a cleaved N-terminal signal peptide (SP), two transmembrane helices (TMHs), and a large extracellular C-terminal domain (10). Both Astn1 and Astn2 undergo an endoproteolytic maturation step in which an unknown protease cleaves the protein just after the second TM segment, with the two fragments remaining attached through a single disulfide bond (10,11).

In the present work, we have mapped the topologies of mouse Astn1 and Astn2 in rough microsomal membranes using glycosylation mapping and protease-protection assays. We find that Astn2 has a cleaved N-terminal SP, an N-terminal domain located in the lumen of the RMs (topologically equivalent to the extracellular surface in cells), two TMHs, and a large C-terminal lumenal domain. We further show that Astn1 has the same topology as Astn2, but see no evidence of SP cleavage for Astn1. Finally, we identify the endoprotease responsible for the maturation of Astn1 and Astn2 as signal peptidase, an ER-localized enzyme that normally removes SPs from secreted and membrane proteins.

## Results

### Predicted topologies of mouse Astn1 and Astn2

Topology predictions for mouse Astn1 (UniProtKB Q61137-1, splicing isoform 1) and Astn2 (UniProtKB Q80Z10-3, splicing isoform 3) produced by the TOPCONS server (12) agree with the topology model for Astn2 derived from epitope tagging and cell-surface staining (11), i.e., an N-terminal signal peptide (SP) followed by two transmembrane segments (TMH1 & 2) and a large C-terminal extracellular domain, Fig. 1. In cells, both Astn1 and Astn2 are cleaved by an unidentified endoprotease into two fragments that remain linked by a disulfide bond (11). Edman sequencing of the two Astn2 fragments showed that the N-terminal one starts at Gly^52^ (just after the predicted signal peptide) and the C-terminal one at Asn^466^ (corresponding to Asn^414^ in the isoform analyzed here). For Astn1, the C-terminal fragment starts at Ser^402^; no sequence could be obtained from the N-terminal fragment in this case.

**Figure 1.**
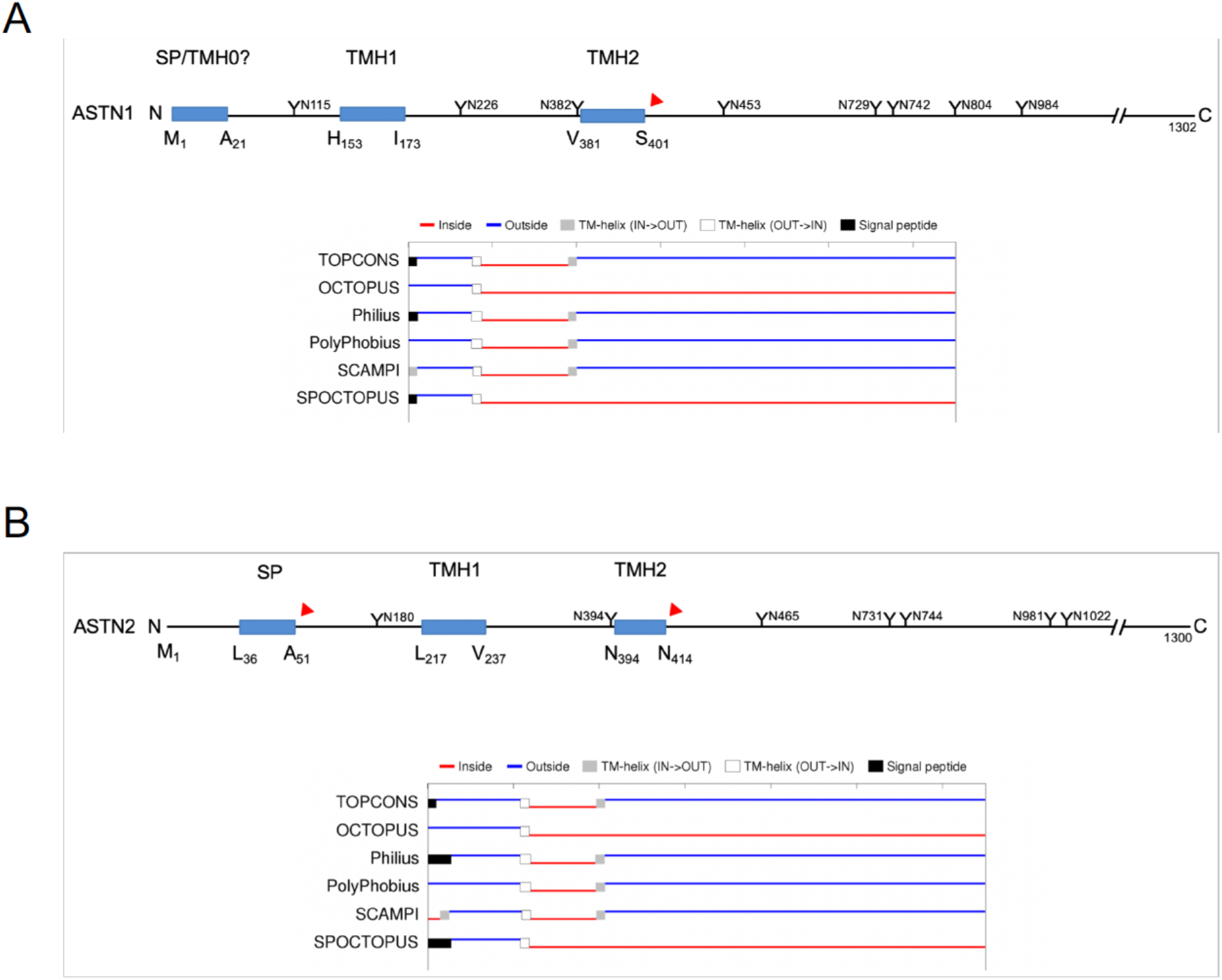
TOPCONS topology predictions. (A) Overview of the sequence of Astn1 with hydrophobic segments (blue), potential acceptor sites for N-linked glycosylation (Y), and proteolytic cleavage sites (red triangles) determined by Edman sequencing (11) marked. The TOPCONS topology prediction (http://topcons.cbr.su.se) is given below. TOPCONS is a consensus predictor that collects data from the other prediction servers listed in the panel. (B) Same for Astn2.

### Topology mapping of mouse Astn1

To characterize the mouse Astn1 protein we used a well-established *in vitro* glycosylation assay (13,14) to determine the topology of the protein when cotranslationally inserted into dog pancreas rough microsomes (RMs). The transfer of oligosaccharides from the oligosaccharide transferase (OST) enzyme to natural or engineered acceptor sites for N-linked glycosylation (-Asn-Xxx-Ser/Thr-Yyy, where Xxx and Yyy cannot be Pro (15-18)) in a nascent polypeptide chain is a characteristic protein modification that can only happen in the lumen of the ER where the active site of the OST is located (19,20). The topology of Astn1 in RMs was also probed by treatment with proteinase K, that can only digest parts of the protein protruding from the cytosolic side of the RMs (21).

To be able to investigate the topology of the 1, 302-residues-long and heavily glycosylated Astn1 protein, we selected to work with various truncated versions of the full-length protein. This was necessary both because *in vitro* translation of such large proteins is inefficient, and because the attachment of an oligosaccharide increases the size of the protein by only 2-3 kDa, a shift that is too small to be detectable by SDS-PAGE for the full-length protein but can easily be visualized when using truncated versions.

Truncated versions of Astn1 were expressed *in vitro* using the TNT^®^ SP6 Quick Coupled System supplemented with column-washed dog pancreas rough microsomes (RMs) (14,21). The glycosylation status was investigated using SDS-PAGE, and truncated Astn1 versions were designed such that differences in glycosylation patterns could be used to infer the topology of the protein in the RM membrane.

As shown in Fig. 2A, Astn1 1-381, a version that extends from the putative SP to the end of the loop between TMH1 and TMH2, receives a single glycan when translated in the presence of RMs (compare lanes 1 and 2). Notably, there is no sign of the SP being cleaved (which would reduce the Mw of the protein by 2.6 kDa). Astn1 78-381 (lanes 3, 4) and Astn1 78-451 (lanes 5, 6) also receive only a single glycan, while Astn1 78-470 (lanes 7, 8) is glycosylated on two sites (note that glycan acceptor sites are rarely if ever modified to 100% in the *in vitro* translation system, hence molecules with both one and two added glycans are visible on the gel). The second glycan addition therefore must be on Asn^453^.

**Figure 2.**
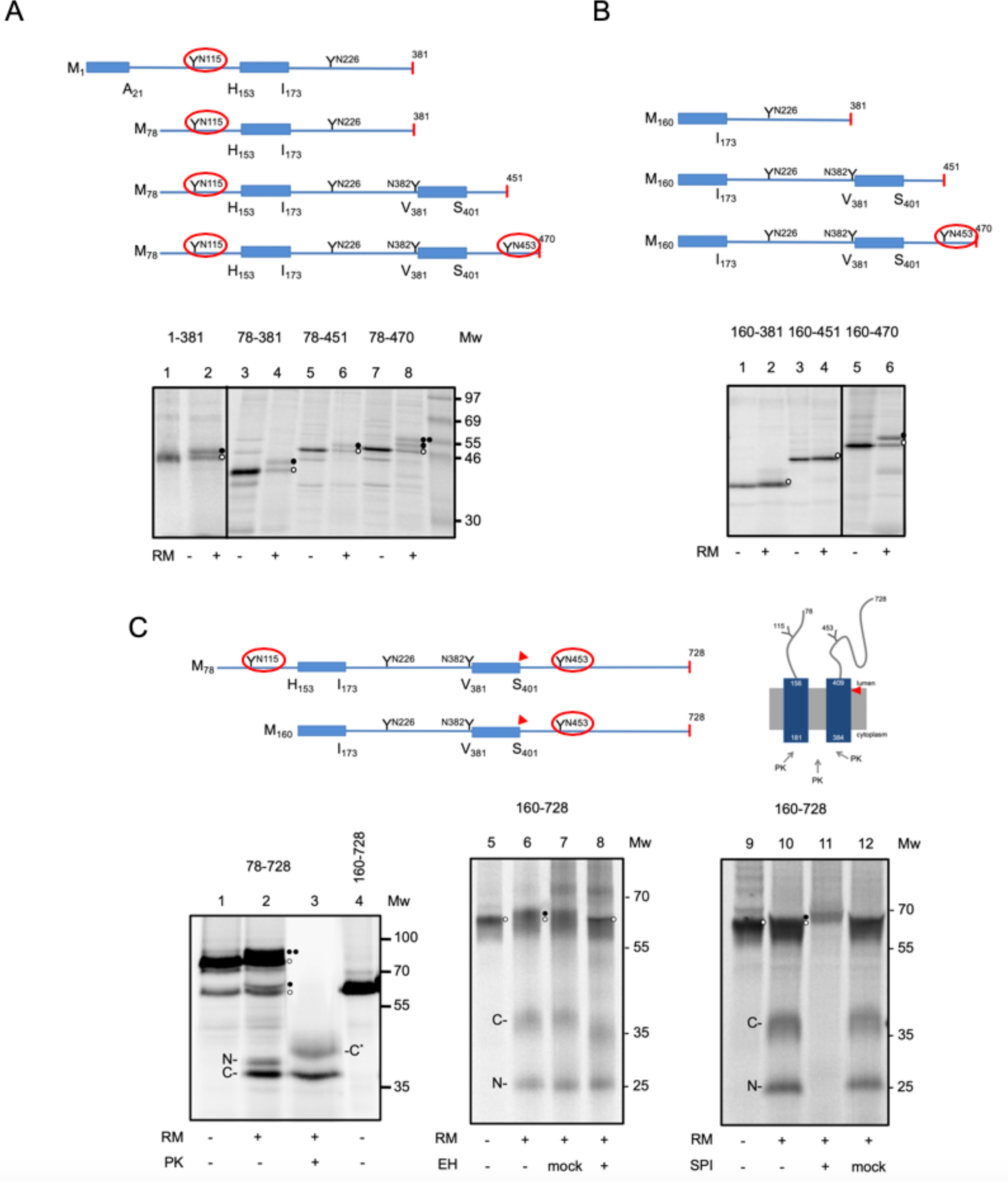
Topology mapping of Astn1 and inhibition of endoproteolytic cleavage by an inhibitor of signal peptidase. (A) The indicated truncated versions of Astn1 were translated *in vitro* with [^35^S]-Met in the presence (+) or absence (-) of RMs, and analyzed under reducing conditions by SDS-PAGE. Unglycosylated products are indicated by an open circle, singly glycosylated products by a filled circle, and doubly glycosylated products by two filled circles. The glycosylated Asn residues are indicated by a red circle in the cartoon. (B) Same as in panel A. (C) Astn1 78-728 was translated *in vitro* with [^35^S]-Met ±RMs (lanes 1 and 2). RMs were subjected to proteinase K (PK) digestion (lane 3). The N- and C-terminal fragments resulting from endoproteolytic cleavage between Ser^401^ and Ser^402^ are indicated (N, C), as is the protease-protected C-terminal fragment (C*). RMs carrying Astn1 160-728 were subjected to EndoH (EH) digestion (lanes 4-8). Note the shift in mobility for the full-length and C bands caused by de-glycosylation (compare lanes 7 and 8). Astn1 160-728 was also translated *in vitro* with [^35^S]-Met in the presence (+) or absence (-) of RMs and the signal peptidase inhibitor N-methoxysuccinyl-Ala-Ala-Pro-Val-chloromethylketone (SPI), lanes 9-11.

To determine whether the first glycan addition is on Asn^115^ or Asn^226^ (Asn^328^ is too close to TMH2 to be reached by the OST (22)), we expressed Astn1 versions lacking the entire N-terminal region, up to but not including TMH2, Fig. 2B. The two shorter versions were not glycosylated at all when expressed in the presence of RMs, while Astn1 160-470 was modified on a single glycosylation site. The latter must be Asn^453^, showing that neither Asn^226^ nor Asn^328^ become glycosylated. We conclude that the putative SP in Astn1 appears not to be cleaved by signal peptidase and probably forms an N-terminal transmembrane helix (TMH0), and that Astn1 has two segments (residues 22-152 and 402-1,302) exposed to the lumen of the RMs, and one segment (residues 174-380) exposed to the cytosol. Further, since Asn^115^ is glycosylated in all four constructs, it appears that the N-terminal segment in the Astn1 constructs that start at M^78^ can be translocated to the lumenal side of the RMs even though it lacks the putative SP.

We further used a protease-protection assay (21) to verify the proposed topology of Astn1. In order that segments of Astn1 that are protected from proteinase digestion by the RM membrane would be of a convenient size for SDS-PAGE separation, we first expressed Astn1 78-728. As seen in Fig. 2C, the protein becomes glycosylated (compare lanes 1 and 2) but it is difficult to determine on how many sites. Interestingly, two prominent bands at ~38 kDa (marked N) and ~36 kDa (marked C) were generated in the presence of RMs (lane 2), suggesting an internal endoproteolytic cleavage, in agreement with the published Edman sequencing results that identified a cleavage site between Ser^401^ and Ser^402^ (1 1). In addition, a third band at ~65 kDa that appears to receive a single glycan in the presence of RMs was also seen (lanes 1 and 2). The latter would be consistent with internal translation initiation at Met^160^, and indeed comigrates with Astn1 160-728 (lane 4).

Proteinase K treatment of RMs carrying Astn1 78-728 digests cytoplasmically accessible parts of the protein and leaves only two protected fragments: one of identical size to the “endoproteolytic” 36 kDa band, and one at ~39 kDa (lane 3). The two protease-protected fragments are precisely what would be expected from the topology derived from the glycosylation study: the 39 kDa band (marked C1*) represents the fragment 381-728 generated when proteinase K digests the cytosolic loop, and the 36 kDa band represents the slightly smaller C-terminal fragment 402-728 generated by endoproteolytic cleavage near the C-terminal end of TMH2. The expected protected N-terminal fragment 78-181 is too small to be resolved on the gel.

Similar results were obtained for Astn1 160-728. In addition to the full-length protein at ~65 kDa, two bands at ~36 kDa (marked C) and ~25 kDa (marked N) were seen in the presence of RMs (compare lanes 5 and 6); EndoH treatment shifted both the full-length band at ~65 kDa and the ~36 kDa band to a lower Mw, while the 25 kDa band did not shift (lane 8). Consistent with the Astn1 160-728 results, the glycosylated ~36 kDa band represents the same endoproteolytic C-terminal fragment 402-728, while the unglycosylated 25 kDa band represents the N-terminal endoproteolytic fragment 160-401.

Given the sequence context of the endoproteolytic cleavage site (see Discussion), we hypothesized that the responsible protease may be signal peptidase. Indeed, inclusion of a signal peptidase inhibitor (23) in the *in vitro* translation of Astn1 160-728 completely inhibits the formation of the ~36 kDa and ~25 kDa products (lane 11).

We conclude that Astn1 has the same topology as previously proposed for Astn2, namely with two lumenal domains (residues 22-152 and 173-1,302) and one cytosolic domain (residues 174-381). The putative SP appears to not to be cleaved, but rather forms an N-terminal transmembrane helix (TMH0). We identify signal peptidase as the enzyme responsible for the endoproteolytic cleavage event at Ser^401^.

### Topology mapping of mouse Astn2

We used the same glycosylation mapping approach to determine the topology of the 1,300 amino acids-long mouse Astn2 protein (splice isoform 3, lacking exon 4 that encodes a 52 residues segment in the domain between TMH1 and TMH2). Astn2 1-482 includes both the putative SP, the two predicted transmembrane helices TMH1 and TMH2, and a portion of the large C-terminal domain. A small amount of glycosylated full-length product at ~56 kDa, two weak bands at ~50 kDa that might represent glycosylated and unglycosylated products lacking the SP (which has a calculated Mw of 6.4 kDa), and a prominent product at ~43 kDa are seen in the presence of RMs, Fig. 3A (lanes 2, 4, 5). The latter is sensitive to EndoH digestion, and the two bands at ~50 kDa collapse to the lower Mw form upon the same treatment (lane 6). The glycosylated 43 kDa band fits the Mw expected for a product resulting from removal of the signal peptide (residues 1-51) and the endoproteolytic cleavage at Asn^413^ observed by Edman sequencing (11) (note that we use a different splice version of Astn2 that lacks 52 residues in the cytosolic segment compared to the one used in this reference). This explains the limited amount of glycosylated full-length product (lanes 2, 4, 5), since most of the molecules that become glycosylated are cleaved after the SP and/or TMH2, as seen in lane 6.

**Figure 3.**
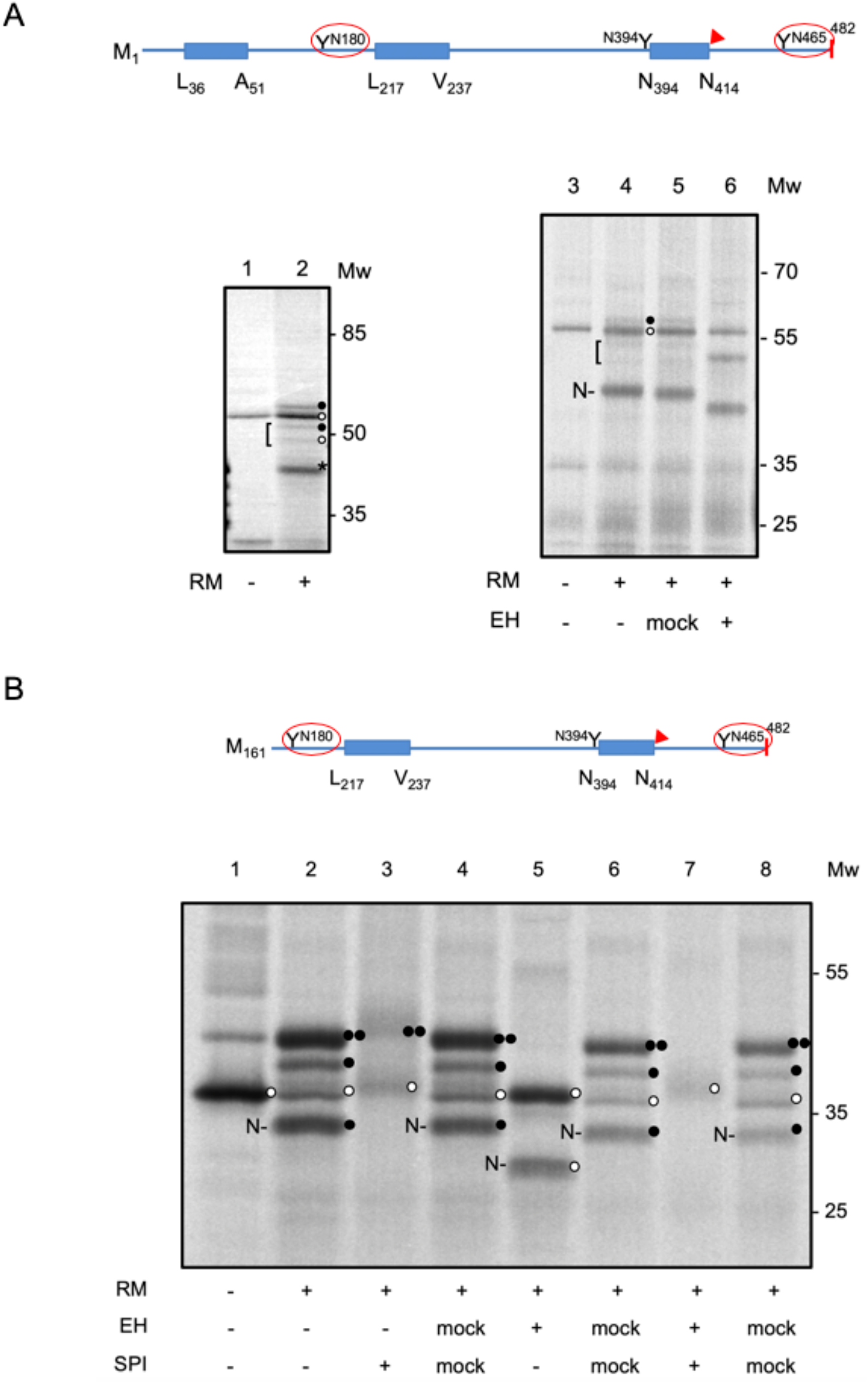
Topology mapping of Astn2 and inhibition of endoproteolytic cleavage by an inhibitor of signal peptidase. (A) Astn2 1-482 was translated *in vitro* with [^35^S]-Met in the presence (+) or absence (-) of RMs, and analyzed under reducing conditions by SDS-PAGE (lanes 1 and 2). Unglycosylated products are indicated by an open circle, and singly glycosylated products by a filled circle. Two cleavage products potentially resulting from removal of the SP by signal peptidase are indicated by a bracket, and the N-terminal endoproteolytic fragment is marked by *. EndoH digestion of RMs with Astn2 1-482 is shown in lanes 3-6; note that the two products potentially generated by removal of the SP (bracket) coalesce into one band and that the endoproteolytic fragment (N) shifts to a lower molecular weight upon de-glycosylation (lane 6). (B) Astn2 161-482 was translated *in vitro* with [^35^S]-Met in the presence (+) or absence (-) of RMs and the signal peptidase inhibitor N-methoxysuccinyl-Ala-Ala-Pro-Val-chloromethylketone (SPI). After translation, RMs were further treated with EndoH (EH) or subjected to mock treatment. The glycosylated Asn residues are indicated by a red circle in the cartoons.

To confirm this interpretation, we also analyzed Astn2 161-482 that lacks the putative SP. As seen in Fig. 3B, Astn2 161-482 yields four prominent bands when expressed in the presence of RMs (lane 2): unglycosylated full-length product at ~37 kDa, singly- and doubly-glycosylated full-length products at ~39 kDa and ~42 kDa, and a smaller endoproteolytic product at ~35 kDa. EndoH treatment collapses the ~39 kDa and ~42 kDa bands to the size of the unmodified full-length product at ~37 kDa, and the ~35 kDa band to a smaller ~30 kDa band (lane 5). Similar to Astn1, addition of the signal peptidase inhibitor to the *in vitro* translation completely inhibits the formation of the ~35 kDa endoproteolytic product (lane 3), and signal peptidase inhibitor plus EndoH treatment of RM-integrated Astn2 161-482 leaves only the unmodified full-length product (lane 7; for unknown reasons, the signal peptidase inhibitor makes bands run slightly higher in the gel).

These results are entirely consistent with the proposed topology of Astn2 (11), and identify signal peptidase as the enzyme responsible for the endoproteolytic cleavage event at Asn^413^.

## Discussion

Earlier work using epitope mapping of Astn2 expressed in COS7 cells have shown that the N- and C-termini are exposed on the cell surface, while the domain between TMH1 and TMH2 can only be immunodecorated in detergent-permeabilized cells (11). Further, both Astn1 and Astn2 were shown to be cleaved by an unknown endoprotease into an N- and a C-terminal fragment, and Edman sequencing of the C-terminal fragments identified cleavage sites between Ser^401^-Ser^402^ in Astn1 and Gly^465^-Asn^466^ in Astn2, just after TMH2. In addition, for Astn2, Edman sequencing of the N-terminal endoproteolytic fragment indicated removal of the putative SP (residues 1-51); no sequence was obtained for Astn1, leaving open whether or not the putative SP is cleaved in this protein.

Here, we have confirmed and extended these results for Astn1 and Astn2 using glycosylation mapping and protease-protection assays in a coupled *in vitro* transcription-translation system supplemented with RMs. Our results for Astn2 are in perfect agreement with those from the earlier study: Astn2 has a cleaved N-terminal SP, an N-terminal domain located in the lumen of the RMs (topologically equivalent to the extracellular surface in cells), two TMHs, and a large C-terminal lumenal domain, Fig. 4. We find that Astn1 has the same topology as Astn2 but see no evidence of SP cleavage; rather, it seems that the putative N-terminal SP in Astn1 remains a part of the protein, presumably forming a third transmembrane helix (TMH0).

**Figure 4.**
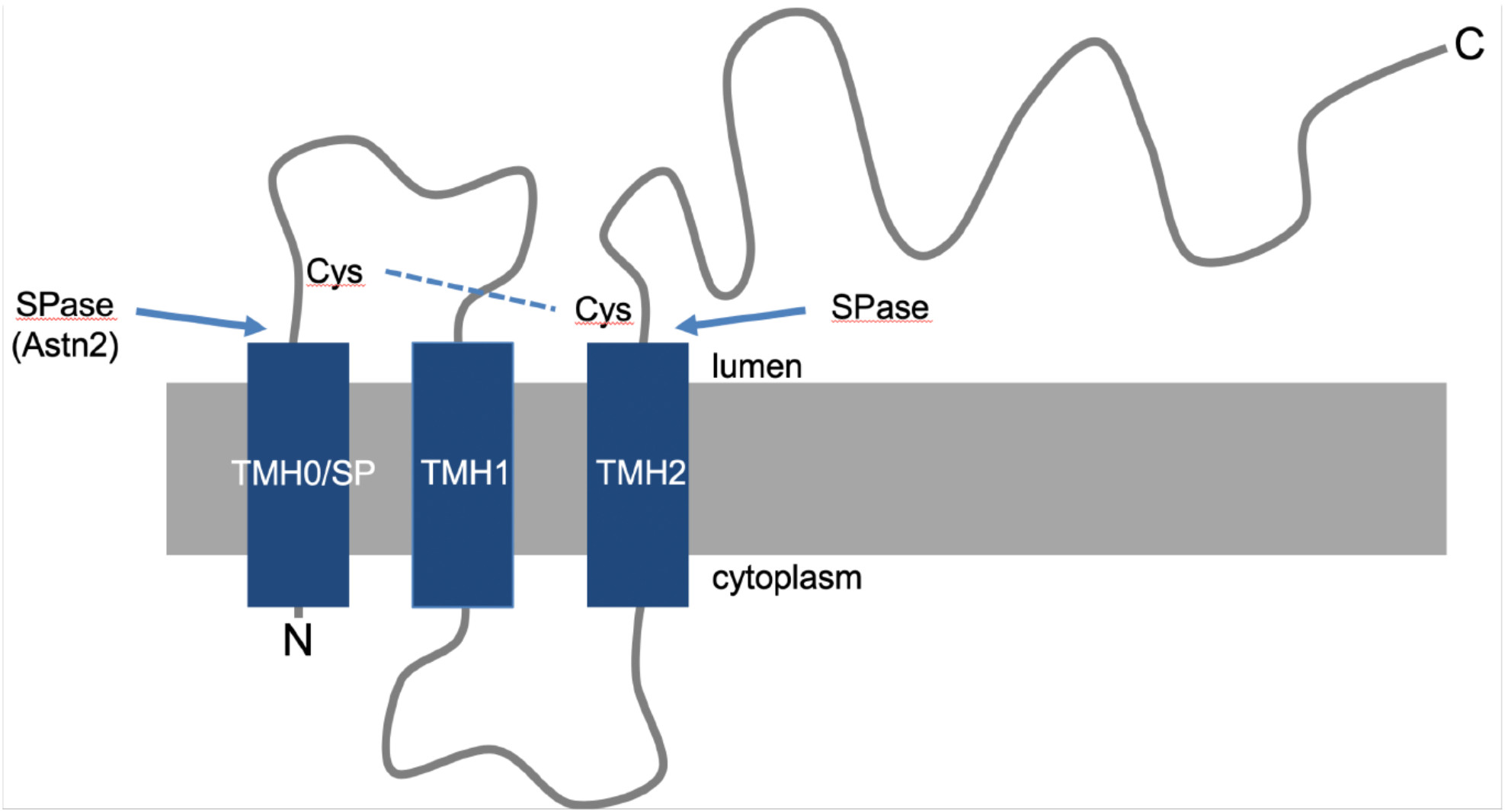
Topology and proteolytic modifications of Astn1 and Astn2. Signal peptidase cleaves both Astn1 and Astn2 after TMH2, and also removes the SP from Astn2. The disulfide bridge that keeps the two endoproteolytic fragments together is indicated.

We further show that an inhibitor of the signal peptidase complex completely inhibits the endoproteolytic cleavage of both Astn1 and Astn2. The unknown endoprotease involved in the maturation of Astn1 and Astn2 is thus signal peptidase, the enzyme that cleaves SPs from secretory and membrane proteins in the ER (24). While it is uncommon that signal peptidase catalyzes internal cleavage reactions of this kind in cellular proteins, many viral polyproteins mature through signal peptidase-catalyzed cleavages after internal hydrophobic segments in the primary translation product (25,26). Indeed, the SP cleavage site and the cleavage site after TMH2 identified by Edman sequencing in Astn2 are precisely the ones predicted by the SignalP server (27), Supplementary Fig. S2.

The present findings raise the possibility that higher levels of SP-mediated cleavage of Astn2 relative to Astn1 explain the higher levels of the Astn1 C-terminus we previously detected on CNS neuronal surface membranes by antibody labeling and functional assays (6,8). This also likely contributes to the apparently distinct functions of Astn1 as a membrane adhesion receptor that functions in glial-guided migration (3,6,7), and of Astn2 as an endolysosomal trafficking protein that functions in both migration (4) and synaptic function (8). Finally, the exceptionally long Astn2 SP hints at the possibility that, after cleavage, the SP may have additional functions in the cell, as seen for many other very long SPs (28). It will therefore be of interest to determine whether the Astn2 SP domain functions in receptor trafficking or planar polarity signaling pathways.

### Experimental procedures

#### Enzymes and chemicals

Unless otherwise stated, all chemicals were from Sigma-Aldrich (St. Louis, MO, US). Plasmid pGEM1, TNT^®^ Quick Coupled transcription/translation system, Rabbit Reticulocyte lysate system and deoxynucleotides were from Promega (Madison, WI, US). [^35^S]-Met was from PerkinElmer (Boston MA, US). All enzymes were from Fermentas (Burlington, Ontario, CA) except Phusion DNA polymerase that was from Finnzymes (Espoo, FI) and SP6 RNA Polymerase from Promega. The QuikChange™ Site-directed Mutagenesis kit was from Stratagene (La Jolla, CA, US) and oligonucleotides were from Eurofins MWG Operon (Ebersberg, DE). All other reagents were of analytical grade and obtained from Merck (Darmdstadt, Germany).

#### DNA manipulations

The cDNAs of mouse astrotactin 1 and 2 (Astn1 and Astn2) (1,302 respectively 1,300 amino acid residues; see Supplementary Figure S1) were cloned into the pRK5 vector using ClaI/SalI (Astn1) and BamHI/XbaI (Astn2) sites. The DNA was then transferred to the pGEMI vector (Promega) at XbaI/SmaI sites together with a preceding Kozak sequence (29), as previously described (13). To create truncations in Astn1, deletions were made between amino acid position 1-78 and 1-160, and stop codons were introduced at positions 382, 452, 471, and 729. Astn2 truncations were created in the same way, with a deletion between 1-161 and a stop codon at 483. The Astn1 and Astn2 cDNAs were amplified by PCR using the *Phusion* DNA polymerase with appropriate primers, and site-specific mutagenesis was performed using the QuikChange™ Site-Directed Mutagenesis Kit from Stratagene. All mutants were confirmed by sequencing of plasmid DNA at Eurofins MWG Operon (Ebersberg, Germany) and BM labbet AB (Furulund, Sweden).

#### In vitro expression

All Astn constructs cloned in the pGEMI and pRK5 were transcribed and translated in an *in vitro* the TNT^®^ SP6 Quick Coupled System from Promega. 150-200 ng DNA template, 1 μl of [^35^S]-Met (5 μCi) and 0.5 μl column-washed dog pancreas rough microsomes (RMs) (tRNA Probes, US) (30) were added to 10 μl of reticulocyte lysate at the start of the reaction, and the samples were incubated for 90 min at 30 °C (21).

#### Proteinase K treatment

PK treatment was performed by adding 1 μl of CaCl2 (200 mM) and 0.2 μl of Proteinase K (4.5 units/ μl) to the translation reaction. After incubating on ice for 30 min, 1 ml of PMSF (20 mM ethanolic solution) was added to inactivate PK and samples were further incubated on ice for 5 min (21).

#### EndoH treatment

For endoglycosidase H (EndoH) treatment 9 μl of the TNT reaction was mixed with 1 μl of 10X glycoprotein denaturing buffer. Following addition of 1 μl of EndoH (500,000 units/ml; NEB, MA, US), 7 μl of d¾O and 2 μl of 10X G3 reaction buffer, and the sample was incubated at 37 °C for 1 h (31). Mock controls were identical, but lacking EndoH.

#### SPI treatment

To demonstrate cleavage by signal peptidase, the inhibitor SPI (N-methoxysuccinyl-Ala-Ala-Pro-Val-chloromethylketone from Sigma) was dissolved in dimethyl sulfoxide (DMSO) and added to the translation mix at a final concentration of 1.4 mM as previously described (14,23,31-33).

#### Analysis and quantitations

Translation products were analyzed under reducing conditions by SDS-polyacrylamide gel electrophoresis, and proteins were visualized in a Fuji FLA 9000 phosphorimager (Fujifilm, Tokyo, JP) using the Image Reader FLA 9000/Image Gauge V 4.23 software (Fujifilm).

## Acknowledgements

We gratefully thank Prof. Arthur E. Johnson, Texas A&M University, for providing dog pancreas microsomes.

## Author contributions

Planned experiments: (PL, ÅT-R, NS, KE, MEH, GvH, IN), performed experiments: (PL, ÅT-R, HB, ZH, NS, KE, MC, AL), analyzed data: (PL, ÅT-R, KE, MEH, GvH, IN), wrote the paper: (MEH, GvH, IN).

## FOOTNOTES

This work was supported by grants from the Knut and Alice Wallenberg Foundation (2012.0282) and the Swedish Research Council (621-2014-3713) to GvH, the Swedish Cancer Foundation (15 0888) to GvH and IMN, the Swedish Foundation for International Cooperation in Research and Higher Education (STINT) (210/083(12); KU 2003-4674) to IMN, the Swedish Foundation for Strategic Research (SSF) (A305:200) and the SSF-Infection Biology (2012(SB12-0026)) to IMN, the Eugene W. Chinery Trust to MEH, and the Renate, Hans, and Maria Hofmann Trust to MEH.

The abbreviations used are:

Astn: Astrotactin
RM: rough microsomes from dog pancreas
OST: oligosaccharyl transferase
ER: endoplasmic reticulum
TM: transmembrane
TMH: transmembrane helix
EndoH: endoglycosidase H
PK: proteinase K
SPI: signal peptidase inhibitor

---

**Supporting Information Figure S1.**
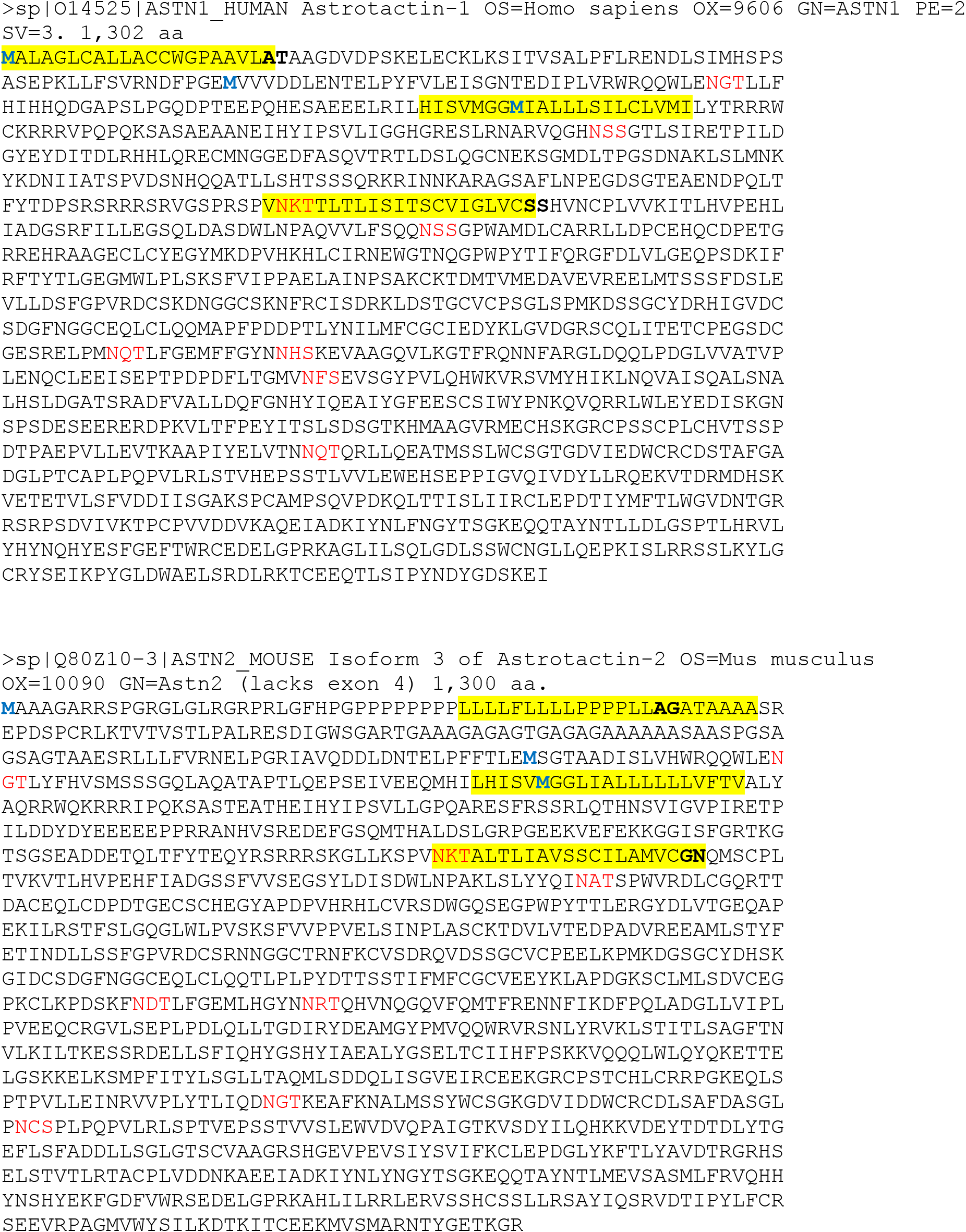
Amino acid sequences of the splice variants of Astn1 and Astn2 used in this study. Hydrophobic regions identified by TOPPRED are shown in yellow, potential acceptor sites for N-linked glycosylation in red, confirmed signal peptidase cleavage sites in bold, and Met residues used as start codons in N-terminally truncated versions in green.

**Supporting Information Figure S2.**
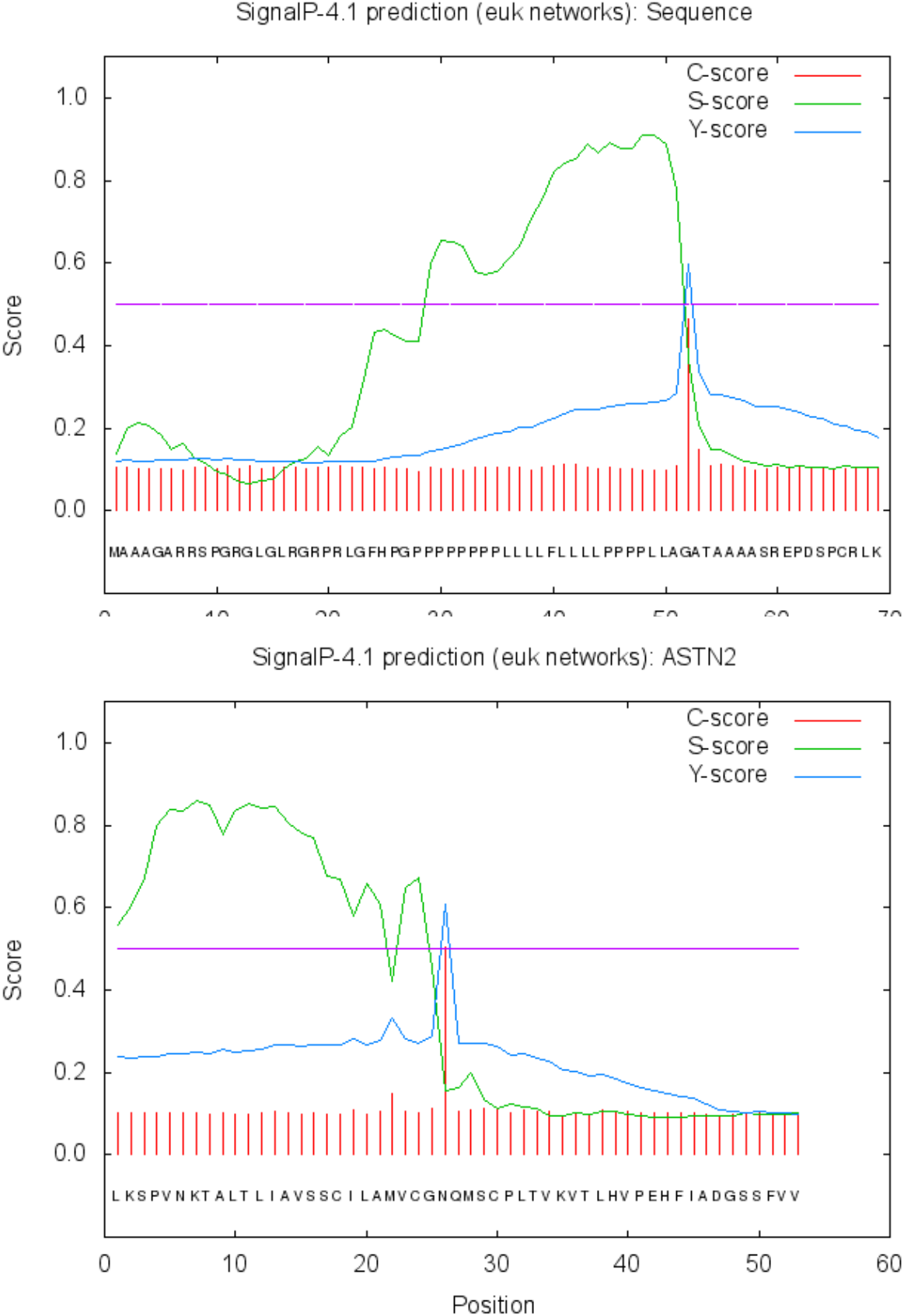
SignalP 4.1 (*Nature methods* 8, 785-786) predicts signal peptidase-catalyzed cleavage (the peak in the C-score) of Astn2 after the SP (Ala51-Gly52; top) and after TMH2 (Gly412-Asn413; bottom). Both sites agree with results obtained by Edman sequencing of the N- and C-terminal fragments o Astn2 (*J Biol Chem* 292, 3506-3516).

## References

1. Zheng, C., Heintz, N., and Hatten, M. E. (1996) CNS gene encoding astrotactin, which supports neuronal migration along glial fibers. Science 272, 417–419

2. Chang, H., Cahill, H., Smallwood, P. M., Wang, Y., and Nathans, J. (2015) Identification of Astrotactin2 as a Genetic Modifier That Regulates the Global Orientation of Mammalian Hair Follicles. PLoS Genet 11, e1005532

3. Edmondson, J. C., Liem, R. K., Kuster, J. E., and Hatten, M. E. (1988) Astrotactin: a novel neuronal cell surface antigen that mediates neuron-astroglial interactions in cerebellar microcultures. J Cell Biol 106, 505–517

4. Wilson, P. M., Fryer, R. H., Fang, Y., and Hatten, M. E. (2010) Astn2, a novel member of the astrotactin gene family, regulates the trafficking of ASTN1 during glial-guided neuronal migration. J Neurosci 30, 8529–8540

5. Ross, M. E., and Walsh, C. A. (2001) Human brain malformations and their lessons for neuronal migration. Annu Rev Neurosci 24, 1041–1070

6. Horn, Z., Behesti, H., and Hatten, M. E. (2018) N-cadherin provides a cis and trans ligand for astrotactin that functions in glial-guided neuronal migration. Proc Natl Acad Sci U S A 115, 10556–10563

7. Adams, N. C., Tomoda, T., Cooper, M., Dietz, G., and Hatten, M. E. (2002) Mice that lack astrotactin have slowed neuronal migration. Development 129, 965–972

8. Behesti, H., Fore, T. R., Wu, P., Horn, Z., Leppert, M., Hull, C., and Hatten, M. E. (2018) ASTN2 modulates synaptic strength by trafficking and degradation of surface proteins. Proc Natl Acad Sci U S A 115, E9717–E9726

9. Ni, T., Harlos, K., and Gilbert, R. (2016) Structure of astrotactin-2: a conserved vertebrate-specific and perforin-like membrane protein involved in neuronal development. Open Biol 6

10. Chang H. (2017) Cleave but not leave: Astrotactin proteins in development and disease. IUBMB Life 69, 572–577

11. Chang, H., Smallwood, P. M., Williams, J., and Nathans, J. (2017) Intramembrane Proteolysis of Astrotactins. J Biol Chem 292, 3506–3516

12. Tsirigos, K. D., Peters, C., Shu, N., Kall, L., and Elofsson, A. (2015) The TOPCONS web server for consensus prediction of membrane protein topology and signal peptides. Nucleic acids research 43, W401–407

13. Lundin, C., Nordstrom, R., Wagner, K., Windpassinger, C., Andersson, H., von Heijne, G., and Nilsson, I. (2006) Membrane topology of the human seipin protein. FEBS Lett 580, 2281–2284

14. Cuviello, F., Tellgren-Roth, A., Lara, P., Ruud Selin, F., Monne, M., Bisaccia, F., Nilsson, I., and Ostuni, A. (2015) Membrane insertion and topology of the amino-terminal domain TMD0 of multidrug-resistance associated protein 6 (MRP6). FEBS Lett 589, 3921–3928

15. Mellquist, J. L., Kasturi, L., Spitalnik, S. L., and Shakin-Eshleman, S. H. (1998) The amino acid following an asn-X-Ser/Thr sequon is an important determinant of N-linked core glycosylation efficiency. Biochemistry 37, 6833–6837

16. Shakin-Eshleman, S. H., Spitalnik, S. L., and Kasturi, L. (1996) The amino acid at the X position of an Asn-X-Ser sequon is an important determinant of N-linked core-glycosylation efficiency. J Biol Chem 271, 6363–6366

17. Igura, M., and Kohda, D. (2011) Quantitative assessment of the preferences for the amino acid residues flanking archaeal N-linked glycosylation sites. Glycobiology 21, 575–583

18. Bano-Polo, M., Baldin, F., Tamborero, S., Marti-Renom, M. A., and Mingarro, I. (2011) N-glycosylation efficiency is determined by the distance to the C-terminus and the amino acid preceding an Asn-Ser-Thr sequon. Protein Sci 20, 179–186

19. Johansson, M., Nilsson, I., and von Heijne, G. (1993) Positively charged amino acids placed next to a signal sequence block protein translocation more efficiently in Escherichia coli than in mammalian microsomes. Mol Gen Genet 239, 251–256

20. Kelleher, D. J., and Gilmore, R. (2006) An evolving view of the eukaryotic oligosaccharyltransferase. Glycobiology 16, 47R–62R

21. Lara, P., Ojemalm, K., Reithinger, J., Holgado, A., Maojun, Y., Hammed, A., Mattle, D., Kim, H., and Nilsson, I. (2017) Refined topology model of the STT3/Stt3 protein subunit of the oligosaccharyltransferase complex. J Biol Chem 292, 11349–11360

22. Nilsson, I., and von Heijne, G. (1993) Determination of the Distance Between the Oligosaccharyltransferase Active Site and the Endoplasmic Reticulum Membrane. J Biol Chem 268, 5798–5801

23. Green, N., Fang, H., Miles, S., and Lively, M. O. (2002) Structure and function of the endoplasmic reticulum signal peptidase complex. in The enzymes (Dalbey, R. E., and Sigman, D. S.eds.), 3rd Ed., Academic press. pp 57–75

24. Nyathi, Y., Wilkinson, B. M., and Pool, M. R. (2013) Co-translational targeting and translocation of proteins to the endoplasmic reticulum. Biochim Biophys Acta 1833, 2392–2402

25. Liljeström, P., and Garoff, H. (1991) Internally Located Cleavable Signal Sequences Direct the Formation of Semliki Forest Virus Membrane Proteins from a Polyprotein Precursor. J Virol 65, 147–154

26. Pene, V., Lemasson, M., Harper, F., Pierron, G., and Rosenberg, A. R. (2017) Role of cleavage at the core-E1 junction of hepatitis C virus polyprotein in viral morphogenesis. PLoS One 12, e0175810

27. Petersen, T. N., Brunak, S., von Heijne, G., and Nielsen, H. (2011) SignalP 4.0: discriminating signal peptides from transmembrane regions. Nature methods 8, 785–786

28. Kapp, K., Schrempf, S., Lemberg, M.K., and Dobberstein, B. (2009) Post-Targeting Functions of Signal Peptides in Protein Transport into the Endoplasmic Reticulum (Zimmermann, R. ed.), Landes Bioscience. 2000-2013. pp 1–11

29. Kozak M. (1989) Context effects and inefficient initiation at non-AUG codons in eucaryotic cell-free translation systems. Mol Cell Biol 9, 5073–5080

30. Walter, P., and Blobel, G. (1983) Preparation of microsomal membranes for cotranslational protein translocation. Methods Enzymol 96, 84–93

31. Lundin, C., Kim, H., Nilsson, I., White, S. H., and von Heijne, G. (2008) Molecular code for protein insertion in the endoplasmic reticulum membrane is similar for N(in)-C(out) and N(out)-C(in) transmembrane helices. Proc Natl Acad Sci U S A 105, 15702–15707

32. Nilsson, I., Johnson, A. E., and von Heijne, G. (2003) How hydrophobic is alanine? J Biol Chem 278, 29389–29393

33. Nilsson, I., Johnson, A. E., and von Heijne, G. (2002) Cleavage of a tail-anchored protein by signal peptidase. FEBS Lett 516, 106–108

